# Urban green infrastructure fails to mitigate exposure to mercury with increasing pollution levels: evidence from corvids

**DOI:** 10.1101/2025.06.13.659450

**Authors:** Michał Ciach, Łukasz J. Binkowski, Arkadiusz Fröhlich, Katarzyna Kucharska

## Abstract

The urban landscape acts as a novel habitat providing opportunities for certain species. However, populations of these so-called urban winners are constantly exposed to environmental contaminants, among which mercury (Hg) is perceived as a significant health concern for both wildlife and humans. While absorption of Hg is primarily related to its environmental level, it could be theoretically mitigated by near-natural habitats that have persisted within the urban landscape. Here, we investigated this hypothesis using three sympatric corvids as a model – Magpie *Pica pica*, Jackdaw *Corvus monedula* and Rook *Corvus frugilegus*. Total Hg concentrations were identified in the feathers (in the shafts and barbs separately) of each species collected in their breeding territories located along the urbanization gradient of Kraków (Poland). These concentrations were then correlated with particulate matter emissions (PM_10_) and habitat features (green and grey infrastructure) measured in the territories where the feathers had been collected. We expected that Hg concentrations in feathers would increase with increasing local pollutant emissions but decrease with increasing areas of green infrastructure, i.e. natural or semi-natural vegetation. Mean Hg concentrations in both feather shafts and barbs differed between the species. Magpie showed the highest concentrations (0.621 ±0.442 SD µg/g in barbs), followed by Jackdaw (0.441 ±0.272 µg/g) and Rook (0.296 ±0.134 µg/g). A generalized linear model yielded a correlation between habitat composition and Hg concentration only for Jackdaw feather shafts. A generalized additive model, however, revealed a non-linear relationship between PM_10_ emissions and Hg concentrations in feather barbs and/or shafts of Jackdaw and Magpie (but not Rook). Hg concentrations initially increased, but then fell with increasing PM_10_ emissions; these relationships were not influenced by habitat features. In conclusion, we found no evidence that Hg contamination was mitigated by urban green infrastructure.

**Highlights:**

- Urban birds are exposed to environmental contaminants, including mercury (Hg)
- Hg concentrations in feather shafts and barbs differed between the species
- Hg concentrations non-linearly correlated with particulate matter emissions
- Hg contaminations not mitigated by urban green infrastructure

## INTRODUCTION

The expansion of urban landscapes in recent centuries has led to their becoming significant parts of terrestrial ecosystems at the global scale (Melchiorri et al. 2018). Built-up areas thus constitute an important habitat for a number of animal species that have successfully adapted to this new type of ecosystem, where they have been able to achieve high population densities (Jokimäki and Suhonen 1998; Mcdonald et al. 2008). Urban-dwelling species benefit from anthropogenic food (Ciach and Fröhlich 2017; Binkowski et al. 2020), mild temperatures (Jadczyk and Drzeniecka-Osiadacz 2013) and reduced natural predation (Malone et al. 2017). However, urban birds are also negatively affected by anthropogenic noise (Nemeth et al. 2013) and artificial lighting at night (Silva et al. 2014). Another factor closely associated with urbanization is environmental contamination (Birke and Rauch 2000; Zhang and Wong 2007; Liu et al. 2010; Binkowski and Meissner 2013). Air, soil and water pollution all expose urban-dwelling animals to various contaminants, including metals (Allison et al. 2006; Rodrigues et al. 2006), which may impair health, reduce reproductive success and lower individual survival rates (Herrera-Dueñas et al. 2014). Such exposure is directly related to the biology of species (Eeva et al. 2009; Berglund et al. 2011), but further bioaccumulation of pollutants and their potential transfer to higher trophic levels also depend on the ecology of species and metal properties (Díez et al. 2008; Orłowski et al. 2010; Binkowski et al. 2024).

One of the most toxic metals is mercury (Hg), which impacts mainly the nervous system, neurological development, the endocrine system and fertility (Scheuhammer 1987; Wolfe et al. 1998; Rutkiewicz et al. 2011; Tartu et al. 2013; Binkowski et al. 2016; Kucharska et al. 2021). The presence of Hg in the environment is linked to volcanism and soil erosion, but also to anthropogenic activities, the main sources being industry, fossil fuel combustion and evaporation by artisanal gold mining (UNEP 2013; Chen et al. 2018). In urban ecosystems, the main sources of Hg include municipal solid waste landfills, waste incineration, consumer products and food leftovers (Muenhor et al. 2009; Tao et al. 2017; Mansouri et al. 2021; EPA, 2025). As Hg is the most efficiently biomagnified of all metals (mainly due to methylation to organic compounds) and in birds is easily deposited into growing feathers, feather concentrations are a good reflection of the exposure of birds to Hg during feather growth (Binkowski et al. 2020; Bottini et al. 2021) and can be used for tracking changes in contamination levels over time (Monteiro and Furness 1997). However, this aspect is also useful when studying spatial differences in contamination levels, e.g. in the urban environment. With such an approach, sedentary urban-dwelling species that spend most of their annual cycle in the urban landscape, where they forage, breed and moult, form a potential model indicator group for environmental pollution and its associated hazards.

Members of the crow family (corvids) *Corvidae* are characterized by large brains, versatile feeding strategies (Kulemeyer 2010) and low levels of neophobia (Greggor et al. 2016). With their high potential to adapt to changing environments, several members of this family have markedly benefited from urbanization, reaching high abundances in urban areas worldwide (Goodwin 1983; Cramp 1994; Anjos et al. 2009). Some corvids, e.g. Rook *Corvus frugilegus*, Jackdaw *Corvus monedula* and Magpie *Pica pica*, have become closely associated with human settlements throughout Europe, and urban landscapes nowadays host a significant fraction of their population during both the breeding and wintering periods (Jokimäki and Suhonen 1998; Ciach and Fröhlich 2017). Corvids are omnivorous, but their feeding preferences depend on the species – from seed eaters and insectivores to predators and scavengers (Holyoak 1968). Moreover, the urban environment provides anthropogenic food, which corvids commonly consume, and may constitute a major fraction of their diet (Marzluff and Neatherlin 2006; Withey and Marzluff 2009). These feeding preferences and the associated differences in dietary composition entail diversified exposure to pollutants in these birds (Díez et al. 2008). As elevated metal concentrations and high levels of dioxins have been detected in urban-dwelling corvids (Benmazouz et al. 2021), this group of birds appears to be an honest indicator of urban pollution.

Given this context, the aim of the present study was to assess the factors influencing Hg contamination in corvids within an urban landscape. We focused on feathers from three sympatric species, i.e. Rook, Jackdaw and Magpie, collected from randomly selected territories in Kraków, Poland, one of the most heavily polluted cities in Europe. We investigated the relationship between air pollution levels (approximated by PM_10_ concentrations) and Hg concentrations in corvid feathers, expecting (1) a positive correlation between environmental pollution and Hg concentrations. As the urban landscape contains a mosaic of habitats with different levels of naturalness, we also investigated the relationship between landcover characteristics and Hg concentrations in feathers. As the presence of habitats consisting of natural vegetation is often regarded as a factor that may limit contamination levels, we anticipated (2) that increasing coverage of green infrastructure, i.e. forests, gardens, parks, farmland and urban greenery within bird territories, would mitigate pollution and be positively associated with reduced Hg concentrations. At the same time, we expected that increasing coverage of grey infrastructure, i.e. built-up areas and roads, would be correlated with Hg concentrations.

## MATERIALS AND METHODS

### Study area

The study was carried out in the city of Kraków, Poland (50°05’ N, 19°55’ E), which covers an area of 327 km^2^ and has a mean population density of 2 331 persons/km^2^ (GUS 2013). Kraków is characterized by a broad urbanization gradient – from the densely built-up city centre, through extensive suburbs with a moderate number of buildings, to the scattered buildings typical of a farmland landscape. Built-up areas cover around 6% of the city’s overall area, and range from the compact, continuous structures covering the ground completely, through taller and shorter blocks of flats to detached and semi-detached houses, with varying amounts of greenery in between. The city’s roads and railway lines make up 4% of its overall area (UMK 2014). Urban greenery covers ∼46% of its overall area and is the predominant form of vegetation in the city; it includes gardens (14%), squares, roadside verges and playgrounds (10%), allotments and orchards (4%), parks and cemeteries (3%) and other green areas (15%). Open areas (37%) include arable land (14%), spontaneous vegetation on fallow land (13%), meadows and pastures (8%) and wetland vegetation (2%). Forests and natural woodland (11%) consist of natural and semi-natural scrub (5%), deciduous and mixed forest (4%), and damp, riparian forest and other tree stands (2%). The quality of air in Kraków is among the worst in Europe, containing high levels of suspended particulate matter, nitrogen dioxide and benzo(alpha)pyrene (WIOŚ 2014; AQIE 2018; Kucharska et al. 2019). In such cities Hg concentrations are thought to be elevated, especially during the winter (Siudek et al. 2015).

### Field methods

At the beginning of bird breeding season in 2014 – 2016, Rook, Jackdaw and Magpie territories were identified along the urbanization gradient from the city centre to the suburbs. Later, at the peak of the moulting season (June and July), nesting sites or foraging grounds were searched for flight feathers. Feathers from Rooks (N = 50), Jackdaws (N = 74) and Magpies (N = 48) were collected and put in a plastic string bag, transported to the laboratory and stored for chemical analysis.

### Analytical methods of Hg quantification

Each feather analysed was washed to remove external contaminants with pure acetone (PA; Roth, GC grade 99.7%) and ultra-pure water (UPW; resistivity of 18.2MΩ cm at 25°C; Direct-Q 3, Merck-Millipore, USA). The washing procedure involved four steps: (1) the feather was placed in a vial with 5 mL PA and agitated on a rotary shaker for 20 minutes; (2) PA was removed and replaced with UPW, after which the vial was agitated on the rotary shaker for another 20 minutes; (3) UPW was removed, the vial with the washed feather was filled with fresh UPW, and the shaking procedure was repeated; (4) the feather was removed from the vial and air-dried at 20-25°C for 24 h. The air-dried feathers were stored in clean test-tubes until analysis.

The total Hg concentration in the feathers was measured using cold-vapour atomic absorption spectrometry (Nippon Instrument Corporation, MA-2) at 253.7 nm. Each feather was divided into two subsamples, the barb and the shaft, from which approximately 6 mg and 9 mg, respectively, were taken for analysis. The samples were not mineralized prior to analysis, but each was supplemented with two additives according to the spectrometer manual, i.e. additives B and M (Wako Pure Chemicals Industries Ltd.), in order to limit potential interferences and further inaccuracies.

The detection limit set for the whole procedure was 0.03 ng per sample. Each sample was measured at least twice, and the mean value was used as the final result expressed as µg of Hg per g of feather (hereafter µg/g). The method was validated against certified reference material (CRM) for human hair ERM-DB001 (JRC, IRMM, Belgium; certified Hg concentration 0.365 ±0.028 mg/g d.w.), which was the closest available matrix to the feathers. The CRM samples were analysed at the beginning and end of each analytical cycle, and their weights were adjusted to represent an amount of Hg similar to that in the feather samples. The measured values for the CRM samples were 0.352 ±0.009 mg/g d.w. (N = 30), showing a recovery of 96.4 ±1.6%.

### Landcover and air pollution variables

The landcover variables were defined within bird territories in a 150 m radius buffer that covered the core part of the species territory (Cramp 1994) on the existing digital spatial database using Geographic Information System tools (QGIS 2013). The total surface areas (ha) of major landcover types of primary importance for the three species were calculated using the polygon vector layer of the atlas of the real vegetation of Kraków (UMK 2014). A separate polygon vector layer was created for each major landcover type. The total surface areas of parks (including parks and cemeteries), gardens (including domestic gardens, allotments and orchards), and forests (including deciduous, coniferous and mixed woodland, and naturally growing shrubs) were calculated. The total surface area of farmland was calculated – this encompassed grasslands (including meadows, pastures, uncultivated and fallow land, swards and heaths) and arable fields (land used for agricultural production). The total surface area of urban greenery represented greenery accessible to the public and included areas situated between buildings and along roadside verges.

Calculation of the built-up areas (ha) was based on the polygon vector layer of terrain covered by compact and continuous buildings, blocks of flats, industrial infrastructure and parking spaces mixed with a fraction of small and highly fragmented green spaces such as lawns, green squares and playgrounds (WODGiK 2015). Calculation of the area of roads and railways (ha) was based on the polygon vector layer of roads and railways (WODGiK 2015), summing the area of infrastructure outlines. Data on emissions of particulate matter of diameter ≤ 10 µm (PM_10_) was obtained from the city’s pollution map (WIOŚ 2014; MIIP 2016), and expressed as the mean of the raster cells falling within the 150 m radius buffer covering the core part of the bird territories. The values taken to construct the variable were the mean annual emissions of PM_10_ particulate matter in the air expressed in kg/year. As PM_10_, PM_2.5_ and other pollutant emissions in the urban environment of Kraków are highly correlated (MIIP 2016), PM_10_ emissions were taken to be a proxy of overall environmental pollution, so that data on other pollutants were not used in the analyses.

### Data handling and analyses

Means ±SD and ranges of Hg concentrations in the shafts and barbs of Rook, Jackdaw and Magpie feathers were calculated and compared between these species using ANOVA tests performed on log-transformed data. A Pearson correlation between Hg concentrations in the shafts and barbs was calculated for each species separately. Means ±SD of habitat variables recorded in the territories of each species were also calculated and compared between the species using ANOVA tests.

Principal component analysis (PCA) was used to construct the index characterizing the habitat composition within the territories of each species. From the variables defining the total surface areas of the major landcover types, i.e. forests, gardens, parks, farmland, urban greenery, built-up areas and roads, the 1^st^ principal component (hereafter – habitat composition index) was obtained, which was used then as an explanatory variable (Fig. 1). The habitat composition index explained 36.2% of the total variance and was an indicator of the urbanization level, as it differentiates sites dominated by roads (PC1 vector and load: 0.55) and built-up areas (0.64) with accompanying urban greenery (0.79) from sites with increasing proportions of wooded habitats, i.e. forests (–0.62), gardens (–0.43), parks (–0.14) and farmland (–0.78) (Fig. 1).

**Fig. 1.**
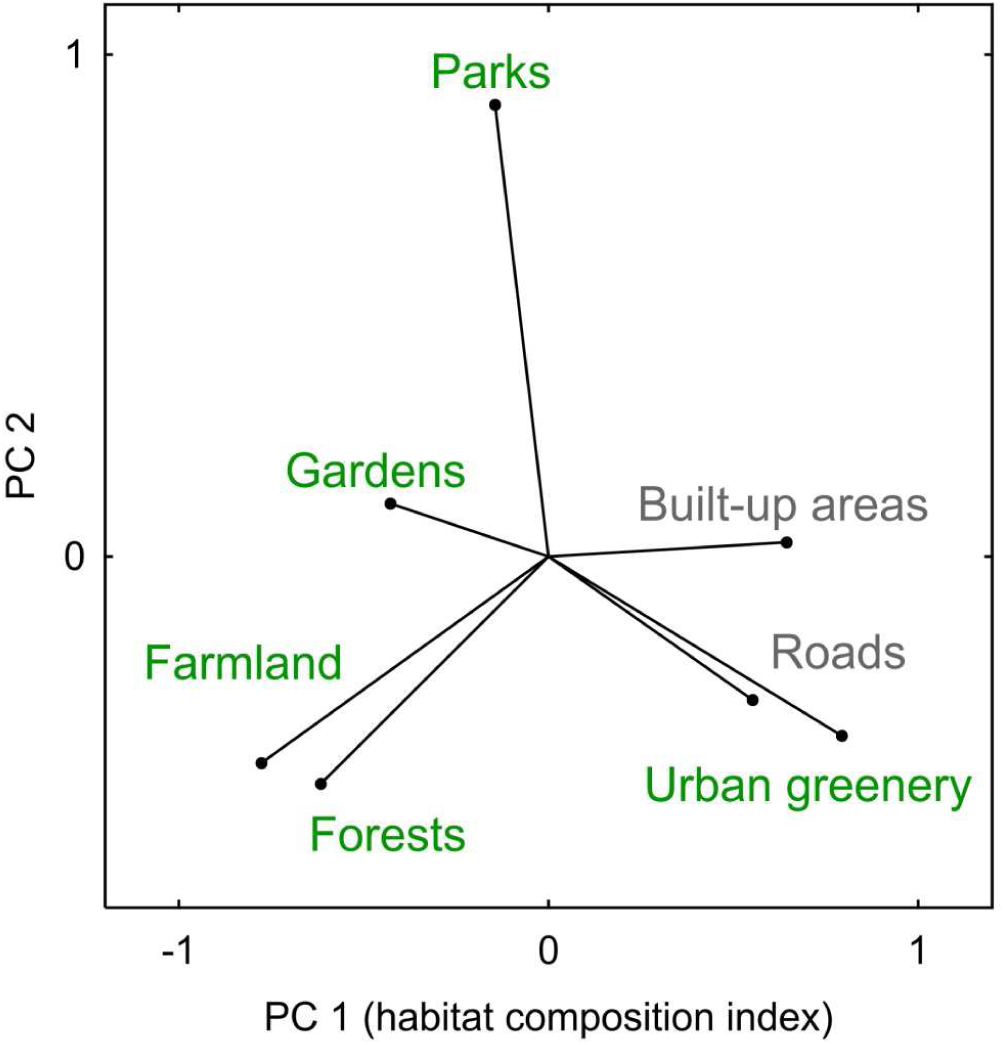
Result of principal component analysis providing vectors and loads for variables contributing to the habitat composition index, i.e. 1^st^ principal component (PC 1), within territories of Rook, Jackdaw and Magpie breeding in the urban landscape of Kraków, S Poland.

Generalized linear models with log-normal error distribution were used to assess the relationship between the habitat composition index, and PM_10_ and Hg concentrations in the feathers of each species. Although generalized linear models imply a linear relationship between the dependent variable and its predictors, we assumed that Hg concentrations in feathers in an urban landscape would also vary in a non-linear manner. Thus, as a further step, we also explored whether Hg concentrations exhibited a potential non-linear dependence on the same variables using generalized additive models (Hastie and Tibshirani 1990; Schimek 2000). A smoothing spline was applied to visualize the revealed relationship between PM_10_ and Hg concentrations in feathers. The statistical procedures were performed using Statistica software (Tibco Inc. 2017). The significance level in all the analyses was set at 0.05.

## RESULTS

All the samples measured contained Hg concentrations above the detection limit. The mean Hg concentration in the feather shafts was 0.108 µg/g, 0.174 µg/g and 0.243 µg/g for Rook, Jackdaw and Magpie, respectively (Table 1) and differed between these species (F_2, 169_ = 23.1, p < 0.001). The mean Hg concentration in the feather barbs was higher than in the shafts for all the species (Table 1) and differed between them (F_2, 169_ = 15.3, p < 0.001), with the lowest values being for Rook (0.296 µg/g), moderate ones for Jackdaw (0.441 µg/g) and the highest ones for Magpie (0.621 µg/g). The correlation between Hg concentrations in the feather shafts and barbs was significant for all three species (p < 0.001). The strongest relationship was noted for Magpie (r = 0.88), followed by the other two species (r = 0.75 in both cases).

**Table 1.** Descriptive statistics of Hg concentrations (µg/g) in feather shafts and barbs of Rook, Jackdaw and Magpie breeding in the urban landscape of Kraków, Poland.

| Species | Feather | Min - max | Median | Geometric mean | Mean ±SD |
| --- | --- | --- | --- | --- | --- |
| Rook<br>(N = 50) | barb | 0.019 – 0.783 | 0.271 | 0.263 | 0.296 ±0.134 |
|  | shaft | 0.010 – 0.305 | 0.100 | 0.092 | 0.108 ±0.058 |
| Jackdaw<br>(N = 74) | barb | 0.103 – 1.267 | 0.329 | 0.369 | 0.441 ±0.272 |
|  | shaft | 0.052 – 0.546 | 0.134 | 0.147 | 0.174 ±0.108 |
| Magpie<br>(N = 48) | barb | 0.166 – 2.114 | 0.468 | 0.510 | 0.621 ±0.442 |
|  | shaft | 0.083 – 0.728 | 0.195 | 0.204 | 0.243 ±0.162 |

The area of landcover types considered as green infrastructure (forests, gardens, parks, farmland and urban greenery) found within the bird territories differed between the species (or approached the adopted significance level) (Table 2). The values of variables regarded as attributes of urbanization, i.e. grey infrastructure (built-up areas and roads), found within the bird territories also differed between the species (Table 2). Although the habitat composition index of the bird territories differed between the species (Table 2), PM_10_ emissions did not display any significant variability between the territories occupied by the three species (Table 2).

**Table 2.** Characteristics of habitat variables analysed in the territories occupied by Rook, Jackdaw and Magpie breeding in the urban landscape of Kraków, Poland, followed by the results of one-way ANOVA.

| Variable | Rook |  | Jackdaw |  | Magpie |  | ANOVA test |  |
| --- | --- | --- | --- | --- | --- | --- | --- | --- |
|  | Mean | SD | Mean | SD | Mean | SD | F<sub>2, 169</sub> | p |
| Forests (ha) | 79.8 | 564.0 | 582.1 | 1896.7 | 1397.5 | 3378.2 | 4.5 | 0.012 |
| Gardens (ha) | 1015.5 | 3753.3 | 2029.1 | 4998.5 | 3573.1 | 6689.0 | 3.0 | 0.054 |
| Parks (ha) | 5149.4 | 7391.8 | 2380.3 | 5568.5 | 3671.1 | 6547.6 | 2.8 | 0.064 |
| Farmland (ha) | 479.8 | 1391.1 | 2087.1 | 4715.4 | 6392.8 | 9969.1 | 12.3 | <0.001 |
| Urban greenery (ha) | 15151.2 | 7277.0 | 14983.2 | 7194.7 | 10214.0 | 9900.9 | 6.2 | 0.002 |
| Built-up area (ha) | 4657.6 | 3400.0 | 5011.1 | 3481.1 | 3358.0 | 2783.9 | 3.8 | 0.023 |
| Roads (ha) | 3281.0 | 2510.1 | 4026.8 | 2400.5 | 1977.2 | 1527.5 | 12.3 | <0.001 |
| Habitat composition index | 0.3 | 0.7 | 0.2 | 0.8 | -0.6 | 1.2 | 15.1 | <0.001 |
| Emission of particulate matter [PM<sub>10</sub>] (kg/year) | 2117.3 | 2322.8 | 2755.7 | 3133.2 | 1778.5 | 1825.2 | 2.2 | 0.110 |

The generalized linear models did not reveal any relationship between the habitat composition index, PM_10_ emissions and Hg concentrations in the feather shafts and barbs of Rook and Magpie (Table 3). In the case of Jackdaw, Hg concentrations in the feather shafts only were negatively correlated with the habitat composition index, indicating that birds occupying territories in areas with increasing coverage of grey infrastructure have higher Hg concentrations (Table 3). The positive relationship between PM_10_ emissions and Hg concentrations in Jackdaw feather shafts approached the adopted significance level (Table 3).

**Table 3.**
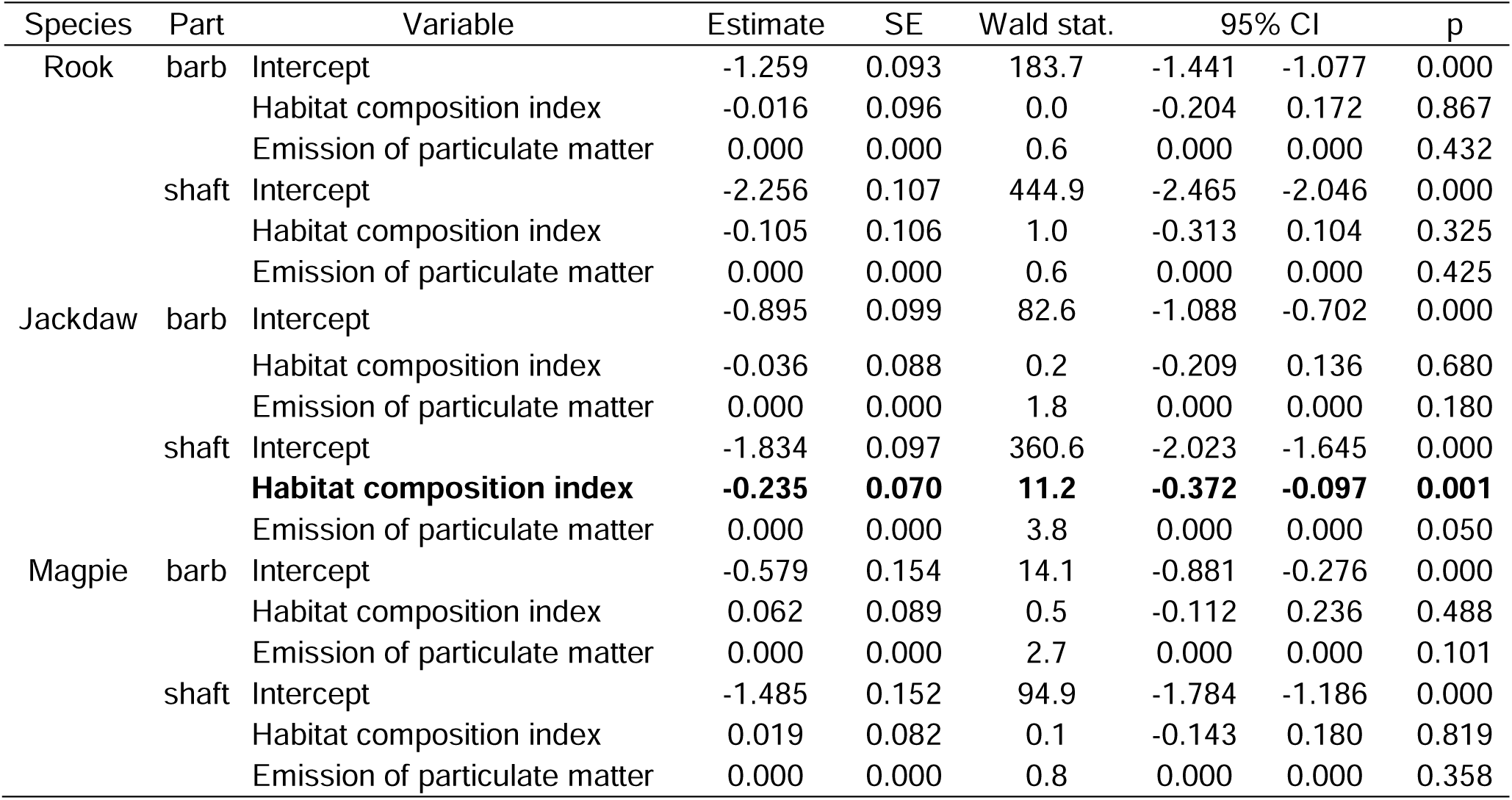
Generalized linear models with estimate, standard error (SE), Wald statistic, 95% confidence interval (CI) and probability (p) illustrating the relationship between the habitat composition index, emissions of particulate matter PM_10_ and Hg concentrations in feathers of Rook, Jackdaw and Magpie breeding in the urban landscape of Kraków, Poland.

The generalized additive models revealed a non-linear relationship between PM_10_ emissions and Hg concentrations in Jackdaw and Magpie feathers (Table 4): Hg concentrations in both species increased with emissions to the level of 5000 kg/year, but then decreased even though emission levels increased further (Fig. 2). Generalized additive models revealed no relationship between Hg concentrations and the habitat composition index in both these species (Table 4). Hg concentrations in Rook feathers were not correlated either with PM_10_ or the habitat composition index (Table 4, Fig. 2).

**Fig. 2.**
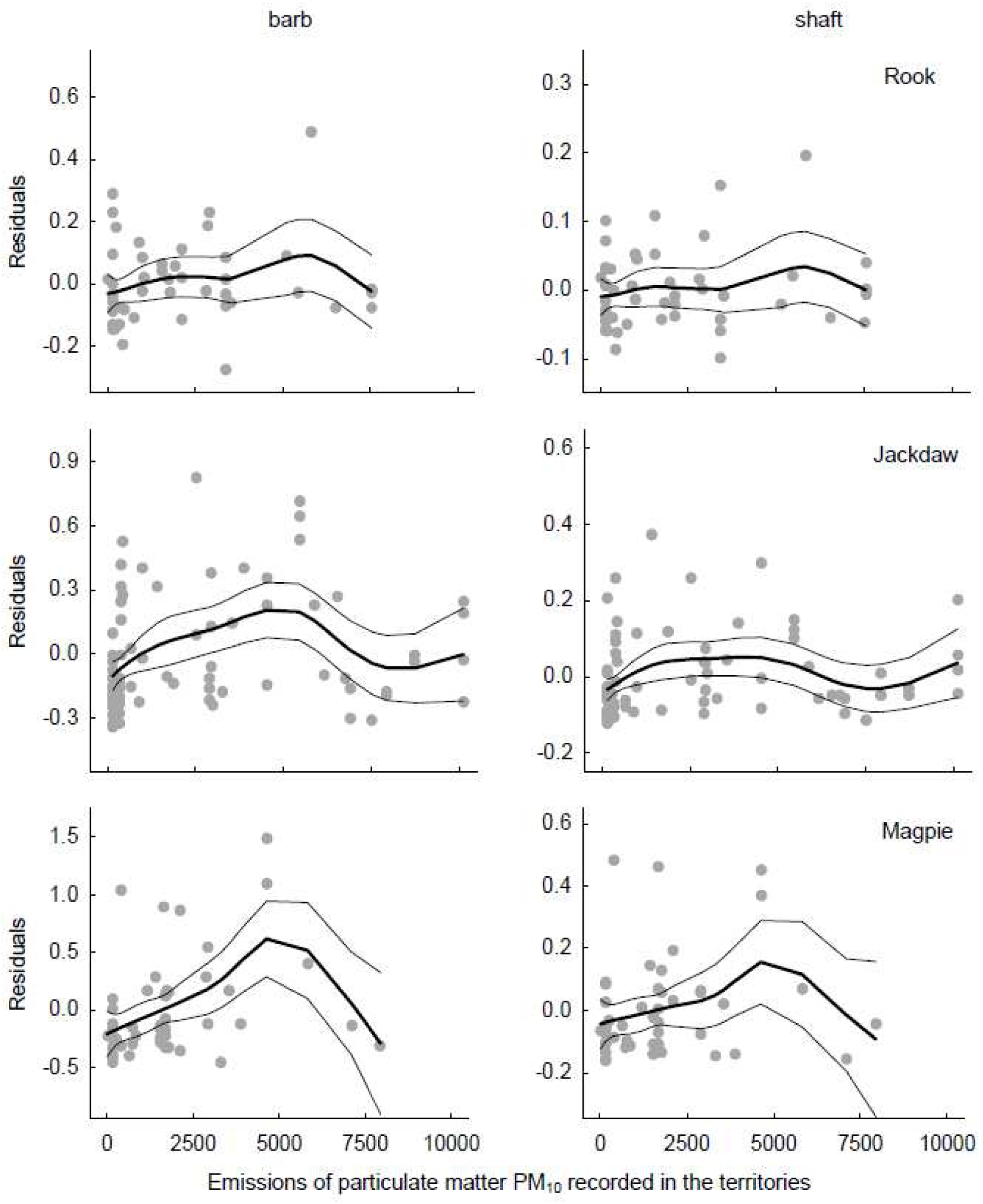
Generalized additive models illustrating the relationship between Hg concentrations in feathers barbs (left-hand column) and shafts (right-hand column) of Rook (upper row), Jackdaw (middle row) and Magpie (bottom row) and emissions of particulate matter PM_10_ recorded in territories located in the urban landscape of Kraków, Poland; bold and thin lines represent the smoothing spline with a 95% confidence interval, respectively.

**Table 4.** Generalized additive models with estimates, standard error (SE) and probability (p) illustrating the relationship between the habitat composition index, emissions of particulate matter PM_10_ and feather Hg concentrations in Rook, Jackdaw and Magpie breeding in the urban landscape of Kraków, Poland.

| Species | Part | Variable | Estimate | SE | p |
| --- | --- | --- | --- | --- | --- |
| Rook | barb | Intercept | 0.290 | 0.028 |  |
|  |  | Habitat composition index | -0.021 | 0.029 | 0.700 |
|  |  | Emission of particulate matter | 0.000 | 0.000 | 0.258 |
|  | shaft | Intercept | 0.108 | 0.012 |  |
|  |  | Habitat composition index | -0.017 | 0.013 | 0.376 |
|  |  | Emission of particulate matter | 0.000 | 0.000 | 0.447 |
| Jackdaw | barb | Intercept | 0.404 | 0.039 |  |
|  |  | Habitat composition index | -0.026 | 0.036 | 0.559 |
|  |  | **Emission of particulate matter** | **0.000** | **0.000** | **0.002** |
|  | shaft | Intercept | 0.164 | 0.015 |  |
|  |  | Habitat composition index | -0.048 | 0.014 | 0.315 |
|  |  | **Emission of particulate matter** | **0.000** | **0.000** | **0.006** |
| Magpie | barb | Intercept | 0.523 | 0.091 |  |
|  |  | Habitat composition index | 0.024 | 0.048 | 0.613 |
|  |  | **Emission of particulate matter** | **0.000** | **0.000** | **0.012** |
|  | shaft | Intercept | 0.223 | 0.037 |  |
|  |  | Habitat composition index | 0.001 | 0.019 | 0.452 |
|  |  | Emission of particulate matter | 0.000 | 0.000 | 0.167 |

## DISCUSSION

### Hg concentrations

We found that Hg concentrations in the feathers of urban-dwelling corvids differed between the feather parts, with higher concentrations in the barbs than the shafts in all three species. According to the literature, feather vanes, consisting of barbs, have higher Hg concentrations than the shafts, which consist of the rachis and calamus (Honda et al. 1985; Peterson et al. 2019). Also, most Hg is concentrated in the feather tip, which consists largely of barbs, and these concentrations are approximately 90% higher than those found in the shaft (Dauwe et al. 2003). The Hg concentration in the shaft makes up 36% of the total feather Hg concentration, while the barb contributes nearly 64% to the feather Hg burden (Peterson et al. 2019). Consequently, removing part of the feather, either barb or shaft, can bias estimates of feather Hg levels (Peterson et al. 2019), resulting in inflated concentrations (Evers et al. 1998; Rimmer et al. 2005; Franceschini et al. 2017). However, in order to track the highest Hg concentrations, it is recommended to check the barb part without the shaft (Solonen and Lodenius 1990; Altmeyer et al. 1991). As many authors analyse the entire feather (or its representative aliquot) without sectioning for different parts, we calculated weighted mean Hg concentrations for whole feathers to facilitate comparison with other studies. Based on the previously mentioned shaft-to-barb Hg concentration ratio (Peterson et al. 2019), the weighted means were 0.23 µg/g for Rook, 0.34 µg/g for Jackdaw and 0.48 µg/g for Magpie.

The Hg feather concentrations found in the three species from Kraków were lower than those measured in Magpie in Italy (1.4-85.9 µg/g; Iemmi et al. 2021) and in other species of birds from around the world (5.6 µg/g; Burger 1993), but were similar to those in various bird species from the Fereydunkenar International Wetland in Iran (0.21 µg/g; Ahmadpour et al. 2016) and in seabirds from Georgia, USA (0.36 µg/g; Becker et al. 2002). The feather Hg levels we found in our study were below the established threshold for sublethal effects, set at 5 µg/g (Eisler 1987; Burger and Gochfeld 1997). However, different avian species may exhibit varying sensitivities to Hg concentrations (Heinz et al. 2009). In some cases, feather Hg levels as high as 40 µg/g have been considered safe as regards reproductive and toxicological effects (Sun et al. 2019), whereas concentrations as low as 0.32 µg/g in other species have been shown to alter antioxidant enzyme activity, leading to lipid peroxidation increase in red blood cells (Espín et al. 2014a, 2014b; Gómez-Ramírez et al. 2023). Therefore, the Jackdaws and Magpies from Kraków, with respective mean whole-feather concentrations of 0.34 and 0.48 µg/g, may be at risk of sublethal effects.

### Feathers as a proxy of Hg exposure

Hg exists in various forms in the environment: elemental (Hg^0^), inorganic (Hg^2+^) and organic (primarily as methylmercury, MeHg) (Yin et al. 2012). In this study we measured the total Hg (THg) concentration, which includes all three forms. The dominant form of Hg incorporated into birds’ feathers is MeHg (Thomson and Furness 1989; Kim et al. 1996; Renedo et al. 2017). Methylmercury tends to accumulate in organisms during their lifetime to much higher concentrations than those in the surrounding environment. Additionally, MeHg is subject to biomagnification, so that animals at higher trophic levels are much more vulnerable (Binkowski et al. 2020). Birds, however, are able to reduce up to 50-93% of their total Hg body burden (Honda et al. 1986; Braune and Gaskin 1987; Lewis and Furness 1991), for example, through depuration into laid eggs and growing feathers. During moulting, MeHg is depurated from the blood into newly grown feathers (Furness et al. 1986; Bearhop et al. 2000; Fournier et al. 2002; Condon and Cristol 2009; Whitney and Cristol 2017), owing to its ability to bind to keratin (Crewther et al. 1965). Importantly, concentrations at the time of feather growth are strongly correlated between feathers and blood, making feathers a good proxy of animal exposure to Hg at that time (Binkowski et al. 2020; Bottini et al. 2021). The complete annual moult of Magpie, Jackdaw and Rook takes place between April and October, but most of the body plumage is moulted in the 3-4 weeks prior to the period when the primaries and secondaries are moulted (Holyoak 1974). All three species start to moult their primaries when their offspring are still in the nest, and the peak of the moulting season in the study area falls between June and August. The moulted feathers from breeding birds were ∼12 months old, while the flight feathers of birds in their 3^rd^ calendar year and older had grown in the previous-year’s moulting season. In contrast, the feathers of birds in their 2^nd^ calendar year grew when they were hatchlings being supplied with food by their parents at the nest.

### Differences between species

We found that Hg concentrations in the feathers of urban-dwelling corvids differ between the species. The highest mean feather Hg concentrations were detected in Magpie, the lowest ones in Rook. The Hg content in feathers is derived mostly from the diet at the time of feather growth (Rogival et al. 2007; Metcheva et al. 2010). During moulting, however, around 70-93% of the internal tissue level of Hg can be remobilized into the feathers, where it binds to the sulphydryl groups in keratin (Honda et al. 1986; Lewis and Furness 1991; Monteiro 1996; Lodenius and Solonen 2013; Renedo et al. 2017), but the rate of Hg depuration from organs to feathers can vary among species (Whitney and Cristol 2017). The differences in feather Hg concentrations between our three species could be due to the compositions of their diets (Anderson et al. 2010; Abbasi et al. 2015; Durkalec et al. 2022). In urban areas, Jackdaw and Magpie feed on a great variety of food, such as plants, seeds, invertebrates and anthropogenic waste (Tatner 1983; Cramp 1994; Meyrier et al. 2017). However, Magpie has a more carnivorous diet than Jackdaw, comprising eggs and nestlings of other species, small vertebrates and their carcasses; it also scavenges roadkill, which has become common in urban ecosystems. Therefore, the diet of a given species may contain more items in which Hg accumulation occurs along trophic chains (Gworek et al. 2020). In the case of Magpie, intensified predation on animals from higher trophic levels that accumulate Hg can lead to such an increase in contamination. Rook, by contrast, relies more on plant-related food sources (Kitowski et al. 2017) and visits rubbish dumps during the breeding season only occasionally, spending more time on foraging outside urban areas, i.e. in farmland (Kasprzykowski 2003). As Hg concentrations in the feathers of granivorous birds tend to be lower than in omnivorous or insectivorous species (Kucharska et al. 2019; Durkalec et al. 2022), the lower Hg concentrations in Rook can be explained by its diet.

### Drivers of Hg concentrations

#### PM_10_ emissions

Our study revealed a positive relationship between PM_10_ emissions and Hg concentrations in Jackdaw feather shafts and barbs and Magpie feather barbs. Hg concentrations in birds increase with the level of urbanization (Pollack et al. 2017; Kucharska et al. 2019), and Hg concentrations in feathers are higher in birds from polluted areas (Di Marzio et al. 2018). Birds can be exposed to Hg through the air (Cizdziel et al. 2013; Cui et al. 2018), although the main route of Hg absorption in birds is via the alimentary tract (Renedo et al. 2018). In the atmosphere, three main forms of Hg occur: particulate bound Hg, gaseous oxidized Hg and gaseous elemental Hg (Selin 2009). The gaseous elemental Hg, which represents >90% of the Hg in the atmosphere, is the most stable form (Fu et al. 2015), can persist for up to 2 years and is the main source of Hg in terrestrial ecosystems (Obrist et al. 2017; Sun et al. 2021). It is mainly inorganic Hg and mono-MeHg deposition that occurs in feathers (Bond and Diamond 2009; Bond and Lavers 2011), hence polluted air may be an immediate source of exposure. Hg concentrations in urban areas are elevated (Siudek et al. 2016) as a result of local emissions, climatic conditions and physical and chemical land coverage (Gworek et al. 2020). Although PM_10_ in Kraków often exceeds permissible values, the concentrations of particulate bound Hg, a constituent of PM_10_, is usually only moderately high, ranging between 0.18-0.49 ng/m^3^ (AQIE 2018; Styszko et al. 2015). The degree to which iHg can influence THg feather concentrations will depend on MeHg concentrations. Assuming that MeHg constitutes approximately 70% of feather Hg concentrations, iHg contributed 0.07, 0.10 and 0.14 µg/g, respectively, of mean THg in Rook, Jackdaw and Magpie, which shows that the impact of iHg increases along with higher MeHg values. Another study has furnished contrasting results: Magpies inhabiting areas distant from urban sites had higher mean Hg concentrations in their feathers than urban Magpies (Iemmi et al. 2021). However, the authors of that study reported a significant variability in Hg concentrations, which will have impacted the mean. They suspected that main explanation for this discrepancy lay in the Magpie’s exploratory behaviour and thus the greater risk of coming into contact with Hg-containing materials while searching for food in urban waste (Iemmi et al. 2021).

#### Influence of habitat

Jackdaws occupying highly urbanized locations had higher Hg concentrations than Magpies and Rooks from less urbanized territories, a pattern which corresponds with other studies (Hargitai et al. 2016; Bauerová et al. 2017). As the concentrations in bird tissues decrease with distance from the source of pollution (Dauwe et al. 2002), habitats located within cities may have different levels of contamination that birds are exposed to (Bauerová et al. 2017). In our study, the territories of each species did not share a common habitat composition pattern but were characterized by different proportions of particular landcover types. In contrast to the two other species, Jackdaws occupied territories with the highest mean proportion of built-up areas. This could have influenced the Hg level because of the higher density of human settlements which contribute to Hg concentrations becoming enriched from the activities of coal combustion facilities, waste incinerators, pharmaceutical waste, food leftovers, paper mills and electrical industries (Sznopek and Goonan 2000; Biester et al. 2002; Tack et al. 2005; Rodrigues et al. 2006).

Jackdaw territories were also characterized by a higher road coverage. A dense road network in urban habitats is a major source of air pollution, which severely reduces individual condition in birds (Esselink et al. 1995; Trombulak and Frissell 2000; Berglund et al. 2011). Higher Hg concentrations from street dust, measured in industrial zones and urban areas (Cai and Li 2019; Dat et al. 2023), can be a source of exposure for the Jackdaws themselves and their prey. Magpie and Rook territories contained more parks, where birds have potential access to feeders and more natural food with a lower Hg load. However, within a given species, our study failed to show a mitigating effect of the increasing proportion of green infrastructure in the bird territory on Hg concentrations. Presumably, gaseous pollutants are easily propagated within the mosaic of urban environments and even larger areas of natural and semi-natural vegetation are unable to sufficiently reduce the spread of Hg-containing substances. Such results indicate the limited role of green infrastructure in mitigating metal contamination in highly urbanized areas, and are an indication of the potential health risks to animals and especially to the humans living there. Although several corvid species are well-known urban dwellers, the influence of pollution on their populations in urban landscapes remains poorly investigated.

### Study limitations

This study offers insight into Hg concentrations in biological samples without the necessity for capturing the birds, a significant advantage. However, from captured live birds, other materials such as blood, fresh faeces and various types of feathers can be sampled, the analysis of which will broaden our understanding of Hg accumulation in a species. Handling live birds can also provide information about aspects of their body condition that may be influenced by Hg exposure.

In our study, we aimed to test the relationships between environmental pollution and exposure to Hg. The PM_10_ parameter we used primarily reflects the inorganic form of Hg, which predominates in the air. As discussed earlier, birds accumulate mainly MeHg, which is primarily associated with the food web. However, MeHg can also be detected in the atmosphere, where it is present as a result of volatilization from sediments, landfills, sewage-sludge, water and other sources containing mercury-methylating bacteria (Carpi et al. 1997; Lindberg et al. 2001; Mester and Sturgeon 2002; Benoit et al. 2003), but the main source is the photodemethylation of volatile dimethylmercury released from landfills (Hintelman 2010). It was shown that atmospheric MeHg is detected mainly in PM_2.5_ (Zhang et al. 2019), so finding a link between PM_10_ and THg concentrations may be challenging. Although there is a distinct correlation between PM_10_ and PM_2.5_ emissions, future research should focus on identifying and incorporating proxies for environmental and trophic MeHg into modelling.

Although we managed to record a non-linear relationship between PM_10_ and Hg concentrations in urban-dwelling corvids, such an association may not be replicable for all species, including humans. The negative effects of Hg contamination are relatively well-recognized in humans (Mergler et al. 2007), as they have been found to contribute to neurodevelopmental disorders (Grandjean and Landrigan 2006), disorders of the endocrine system (Tan et al. 2009), and coronary heart and cardiovascular diseases (Virtanen et al. 2005; Houston 2011). The results of our work, indicating the lack of a mitigating effect of green infrastructure on Hg concentration, have important implications for the health of urban residents. However, the relationship between pollutant emissions in urban landscapes, urban greenery and the levels of contamination in humans is difficult to assess.

## CONCLUSIONS

The study revealed a non-linear relationship between Hg concentrations in the feathers of urban-dwelling corvids and pollutant emissions. Increasing levels of environmental pollution at the local scale lead to higher Hg concentrations in tissues up to a certain point, but these then decrease, even though pollutant emission levels continue to rise. Hg concentrations were not mitigated by the availability of natural and semi-natural vegetation, which indicates the limited role of green infrastructure in counteracting the impact of pollution on organisms inhabiting urban landscapes. The flight feathers of urban-dwelling corvids constitute valuable research material, as they can be collected non-lethally, without the need to capture and stress the birds. Feathers of species with a known moulting pattern, like corvids, may act as an indicator of local environmental Hg exposure (including the trophic circulation) in the absence of other biological samples.

## Acknowledgments

This study was financially supported by the Ministry of Science and Higher Education of the Republic of Poland within the framework of the statutory (DS-3421/2017-2018) and subsidy (2019-2020) funds awarded to the Faculty of Forestry, University of Agriculture in Kraków and statutory subsidy (BS-458/G/2018) funds awarded to the Institute of Biology and Earth Science, University of the National Education Commission, Kraków.

## Ethical statement

The study was performed in accordance with Polish law.

